# Missense *LMNA* Variant Compromises Nuclear Integrity and Sarcomeric Remodeling in Dilated Cardiomyopathy

**DOI:** 10.64898/2026.09.21.753175

**Authors:** Tanushri Dargar, Estèle Lafont, Laura Boulogne, Ludovic Gomez, Maïté Carre-Pierrat, Vanessa Del Vitto, Anna Rausch De Traubenberg, Alexandre Janin, Gilles Millat, Anne Gaignerie, Aude Derevier, Marie Abitbol, Philippe Chevalier, Vincent Gache

## Abstract

Dilated cardiomyopathy (DCM) is a leading cause of heart failure and cardiac transplantation, and pathogenic variants in *LMNA* are a well-established cause of inherited DCM. The *LMNA* gene encodes nuclear lamins A/C, which maintain nuclear integrity, regulate gene expression and mediate mechanotransduction. Here, we investigated the pathogenic consequences of the NM_170707.4(*LMNA*):c.274C>T NP_733821.1:p.(Leu92Phe) variant, previously associated with lipodystrophy features, using patient-derived induced pluripotent stem cells, differentiated into cardiomyocytes and show implication of *LMNA* in sarcomere remodeling and mitochondria efficiency.

We generated iPSC lines from two DCM patients carrying *LMNA* p.Leu92Phe variant in heterozygous form and a healthy parental control. Cardiomyocytes differentiation efficiency was preserved, however, *LMNA* p.Leu92Phe iPSC-CMs exhibited laminopathies associated phenotypes, such as nuclear shape abnormalities and lamin A/C aggregation. Moreover, *in vitro* study revealed that *LMNA* p.Leu92Phe iPSC-CMs alter sarcomere reformation and decrease mitochondrial respiration after cardiomyocyte remodeling, which is associated with a worsening nuclear shape phenotype. Functional analyses highlight defects in calcium handling, thereby explaining arrhythmia and dilated cardiomyopathy features in patients.

Our results show that the *LMNA* p.Leu92Phe variant compromises nuclear lamina integrity and disrupts functional cardiomyocyte properties, particularly during sarcomere remodeling, highlighting the long-term impact of this specific variant in *LMNA*-associated DCM.

**Author Summary:** Specific genetic change in the LMNA gene causes serious heart condition called dilated cardiomyopathy. This condition weakens the heart muscle and can lead to heart failure. We used stem cells from patients carrying a specific mutation and turned them into heart cells to identify altered mechanisms.

We found that even though these cells developed normally at first, they showed clear problems once they matured. The nuclei inside the cells became misshapen, and the structural proteins clumped together abnormally. More importantly, we discovered that the mutation disrupted how heart muscle fibers rebuild themselves and reduced the energy production in mitochondria. We also observed problems with how the cells handle calcium, which could explain why patients experience irregular heartbeats.

Our work shows that this particular genetic variant damage the nuclear structure and impairs critical heart cell functions, especially during muscle remodeling. This helps explain why people with this mutation develop progressive heart disease over time.

## Introduction

Dilated cardiomyopathy (DCM) is a leading cause of heart failure and pathogenic variants in *LMNA* [MIM:150330]^1^are well-established cause of inherited DCM^2^. Lamins A/C, encoded by the *LMNA* gene, are intermediate filament proteins that assemble to form the nuclear lamina contributing to nucleus structural integrity, chromatin organization and genome expression^3^. Laminopathies present nuclear envelope shape deformation/rupture and defective mechanotransduction associated with higher arrhythmic risk^3-8^. Among *LMNA* variants, the NM_170707.4(*LMNA*):c.274C>T NP_733821.1:p.(Leu92Phe) variant, located on the α-helical rod domain, was previously identified in a genetic screen for DCM^6^ and reported in association with Familial Partial Lipodystrophy Type-2 (FPLD2)^7^.

In this study, we employed iPSC-derived cardiomyocytes from two relatives carrying the *LMNA* p.Leu92Phe variant in heterozygous form, which manifests predominantly cardiac phenotypes rather than lipodystrophy features. We show that *LMNA* p.Leu92Phe variant does not affect cardiomyogenesis but leads to alterations in nuclear morphology, alters sarcomerogenesis remodeling and impairs mitochondrial metabolic efficiency, ultimately contributing to an arrhythmic phenotype, confirming the pathogenesis of the *LMNA* p.Leu92Phe variant.

## Results

### Patients with Dilated Cardiomyopathy harbor *LMNA* p.Leu92Phe variant in heterozygous form

In 2010, Hospital Civil of Lyon (HCL) recruited a family pedigree including the proband (III-4) and his daughter (IV-2) (Fig. 1a). Proband was initially diagnosed with arrhythmia and dilated cardiomyopathy^6^. Both later underwent heart transplantation (Table S1). To establish an accurate genetic diagnosis, we performed genetic testing and identified a missense variant in the *LMNA* gene [*c*.*274C>T*; p.(Leu92Phe)] in both affected individuals. *In silico* analyses show highly conserved residues in the rod domain region of lamin across species (Fig. S1a-b). This variant was classified as likely pathogenic using the American College of Medical Genetics and Genomics (ACMG) guidelines using the criteria (PP1_moderate, PP3, PP4, PM2, PS4_moderate)^9^. Combined with clinical features, both patients were diagnosed with *LMNA*-associated dilated cardiomyopathy (*LMNA*-DCM).

**Figure 1:**
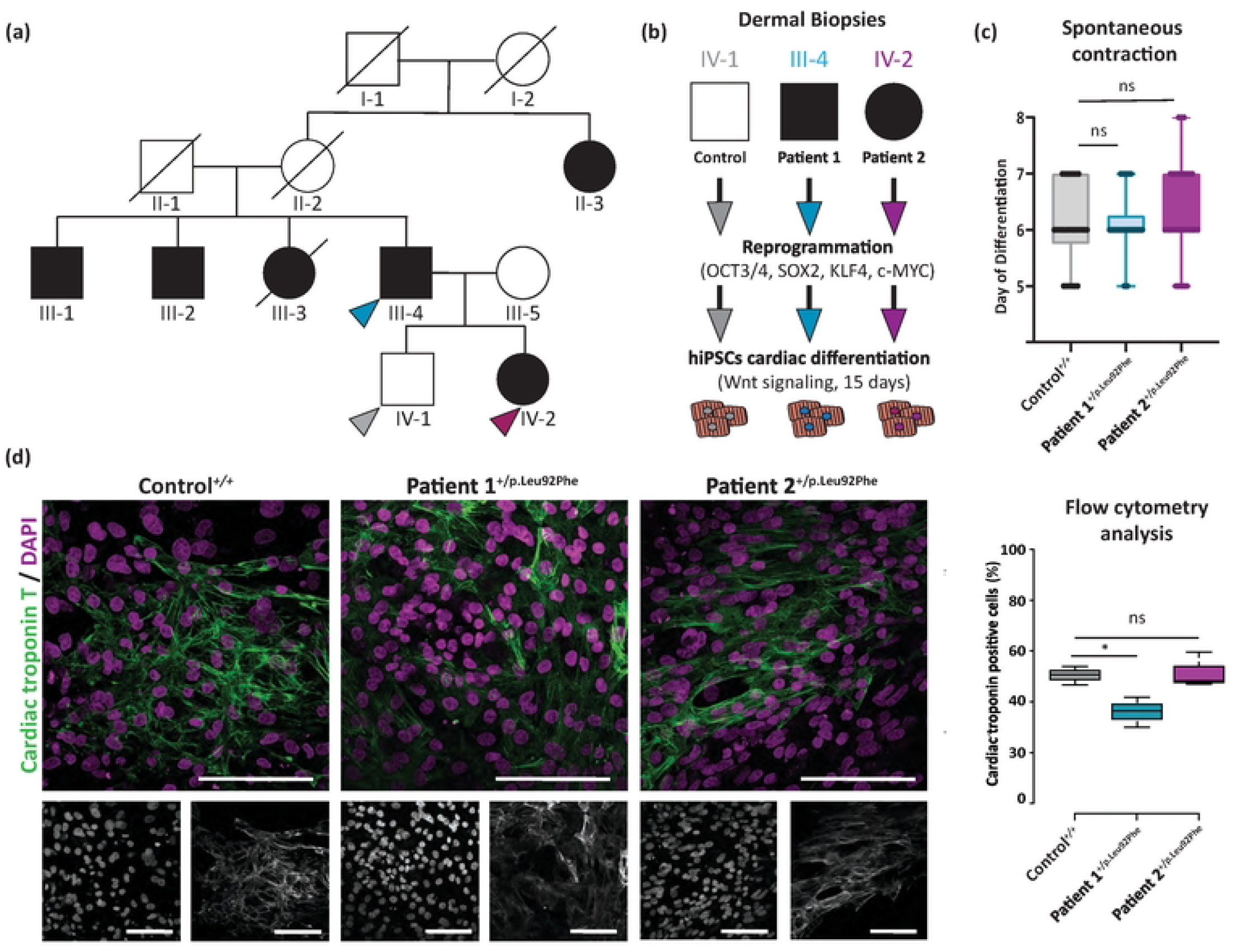
Cardiac differentiation of iPSC derived from DCM-affected individuals harboring the LMNA:p.Leu92Phe variant in heterozygous form. (a) Family pedigree of the proband (III-4) showing autosomal dominant inheritance of the dilated cardiomyopathy and correlation with the *LMNA* p.Leu92Phe variant. The proband (Patient 1, III-4), Patient 2 (IV-2), and an unaffected member used as Control (IV-1) are indicated by green, magenta, and grey arrowheads, respectively. Dilated cardiomyopathy-affected individuals are depicted in black. Unaffected individuals are represented with empty symbols. Deceased individuals are barred. The two deceased grandparents (I-1 and I-2) were not genotyped. (b) Schematic representation of the pipeline to obtain patient iPSC derived cardiomyocytes. (c) Graph representing the initiation of contraction after the beginning of cardiac differentiation (N=30 independent differentiations); Mann-Whitney test for non-normal distributions was performed between control and patients individually. The graph represents the minimum and maximum values with a line at the median. (d) Representative images illustrating typical iPSC derived cardiomyocytes 15 days post differentiation, labelled with cardiac troponin T in green and nuclei with DAPI in magenta. Scale bar: 100µm. Flow cytometry quantification of the total percentage of cardiac troponin-positive cells across lines, utilized to assess overall differentiation purity after 15 days in differentiation. Normality was assessed using the Shapiro-Wilk test. The groups were compared using Mann-Whitney test; the p-value is represented as <0.0332 (^*^) (N=3 independent differentiations).

### *LMNA*-variant iPSCs preserve cardiac differentiation potential

To explore the functional consequences of the *LMNA* p.Leu92Phe variant, we generated patients’ induced pluripotent stem cell (iPSC) lines (Fig. 1b, S2 and Table S3), differentiated into contractile cardiomyocytes (iPSC-CMs)^10^ (Fig. 1b). Patients and control iPSC-CMs initiated spontaneous contractions after 6 days, indicating that the *LMNA* p.Leu92Phe variant does not impair early cardiac specification (Fig. 1c).

Immunostaining using cardiac troponin T (cTnT) confirms cardiac differentiation but reveals variable potency as not all cells were positive for cTnT (Fig. 1d). Flow cytometry analysis of cardiomyocytes showed that in our conditions, nearly 50% of differentiated cells express cTnT with slight patient’s variation (Fig. 1d, Fig. S3), representing standard differentiation heterogeneity after reprogramming^11^. These findings show that cardiac specification is preserved with *LMNA*p.Leu92Phe variant.

### *LMNA*-variant iPSC-CMs display lamin A/C aggregates and nuclear shape abnormalities

To further confirm laminopathy-related phenotypes, we examined nuclear architecture and lamin A/C organization in cardiomyocytes. To reduce cellular heterogeneity, cardiomyocyte population was enriched using lactate-based metabolic treatment^12^. Immunostaining of lamin A/C showed that the protein is still present inside nuclei, suggesting that the *LMNA*p.Leu92Phe variant did not impact protein production or nuclear lamina assembly (Fig. 2a). Quantitative analysis of nuclear cross-sectional area and minimal nuclear Feret’s diameter shows a significant reduction in patient-2 iPSC-CMs compared to control, whereas patient-1 displayed values comparable to the control (Fig. 2b). A key event which links *LMNA* mutations is lamin aggregation and lamina imbalance, reflecting a breakdown in nuclear lamina homeostasis^13^. In control iPSC-CMs, lamin A/C was uniformly distributed along the nuclear envelope. By contrast, both patient-1 and -2 derived iPSC-CMs exhibited a 3-fold increase in the proportion of focal lamin A/C aggregates (Fig. 2c). Similarly, lamin A/C and Lamin B imbalance, that reflect the nuclear rim continuity is increased by 4- and 5-fold in patient-1 and -2 respectively (Fig. 2d). These findings indicate that *LMNA* p.Leu92Phe variant promotes lamin A/C aggregation, perturbing lamin B1 organization.

**Figure 2:**
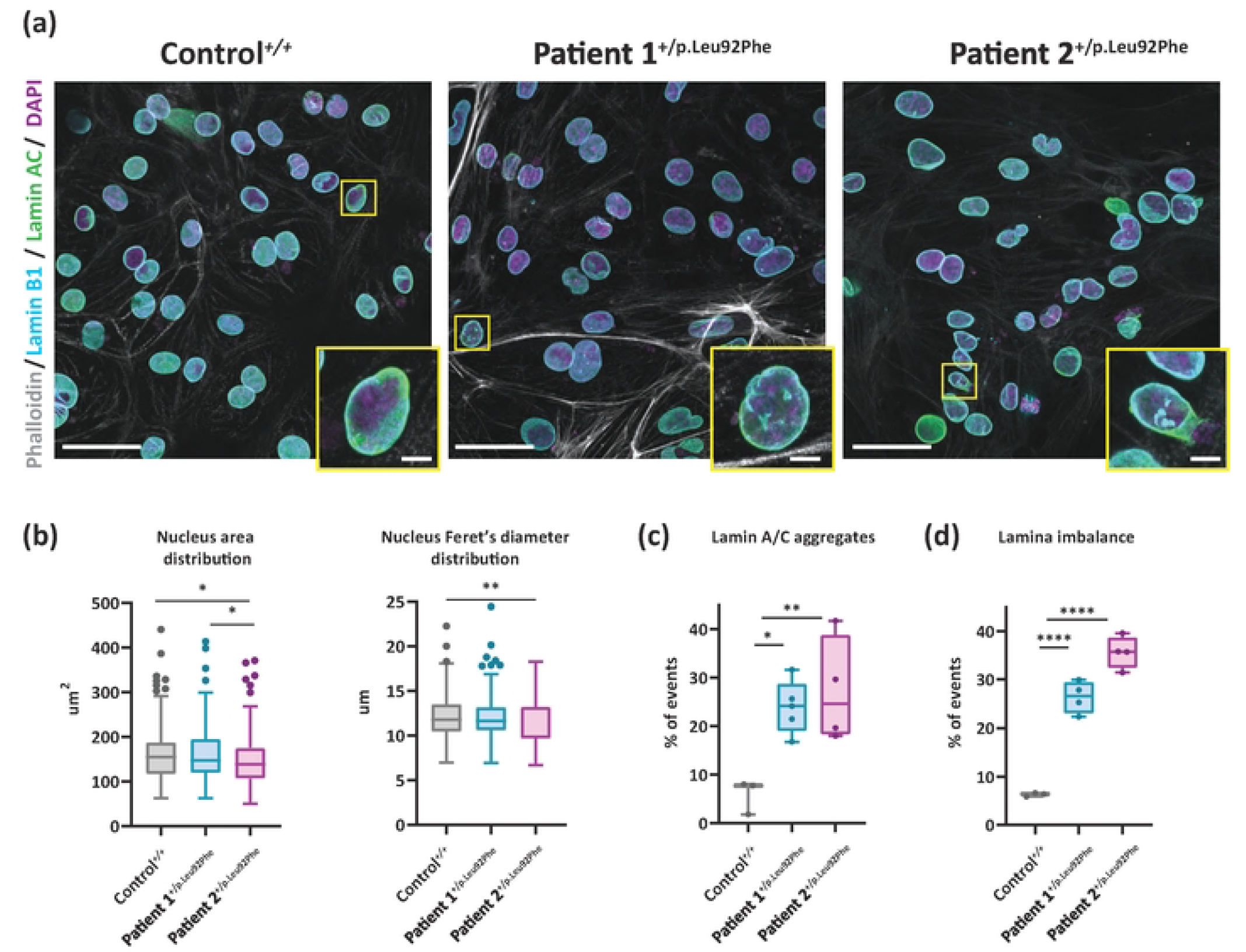
*LMNA*-variant iPSC-CMs exhibit lamin A/C aggregates, nuclear shape abnormalities. (a) Representative images depicting the hiPSC-derived cardiomyocytes after day 25, labelled with lamin A/C in green, lamin B1 in cyan, phalloidin/actin in grey and nuclei with DAPI in magenta. Scale bar: 50µm; Nuclei with lamina imbalance and lamin A/C aggregates are zoomed in, Scale bar: 5µm (b) Quantification representing nuclei area and nuclei Feret’s diameter. N=3 (∼50 nuclei per replicate). Mann-Whitney test to compare patient and control groups. p-value shown as 0.123 (ns), 0.0332 (*), 0.0021 (**). The graph represents Tukey’s plot with a line at the median. (c) Quantification of lamin A/C aggregates in the patient iPSC-CMs as compared to the control. Control, N=3; patient 1 and patient 2, N=4, Ordinary one-way analysis of variance (ANOVA) followed by Tukey’s multiple comparison test. p value shown as 0.0332 (*), 0.0021 (**). The graph represents a box plot, with the minimum and maximum values indicated by lines, and the median marked by a line at the center. (d) Quantification of lamina imbalance in the patient iPSC-CMs as compared to the control. Control, N=3; patient 1 and patient 2, N=4, Ordinary one-way analysis of variance (ANOVA) followed by Tukey’s multiple comparison test. p-value shown as <0.0001 (^****^). The graph represents a box plot, with the minimum and maximum values indicated by lines, and the median marked by a line at the center.

### *LMNA*-variant iPSC-CMs exhibit altered sarcomere reformation, correlated with nuclear shape abnormalities

DCMs are mainly associated to impairment in sarcomerogenesis^14^. We thus applied a “Sarcomeric Remodeling” protocol (SR) to decipher cardiomyocyte’s ability to perform sarcomere reformation (Fig. 3a). iPSC-CMs were enzymatically detached and replated on Matrigel® coated support for 5 days to triggers partial sarcomere disassembly^15^ and provide a assay to monitor sarcomere reassembly (Fig. S3b). After SR protocol, in control conditions, 87 ± 8 % of iPSC-CMs are positive for cardiac troponin and up to 93 ± 0.1 % successfully reformed striated sarcomeres (Fig. 3a-b). By contrast, patient 1- and -2 iPSC-CMs showed only 59 ± 14 % and 47 ± 11 % cardiac troponin positive cells, respectively, and the majority of cardiomyocytes failed to re-establish sarcomeres (Fig. 3a-b, S3c).

**Figure 3:**
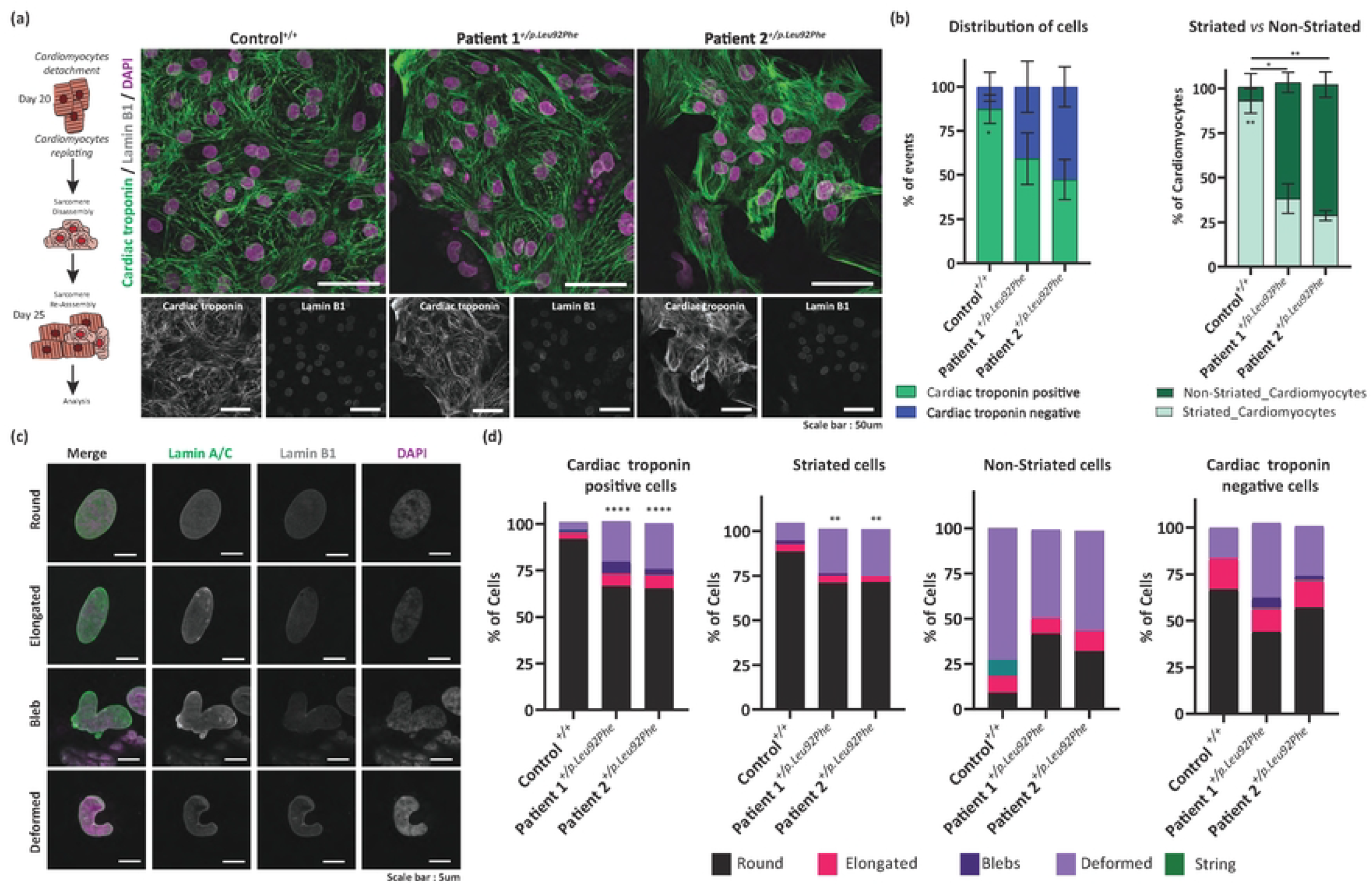
LMNA variant iPSC-CMs exhibit impaired sarcomere reformation with increased nuclear shape abnormalities. (a) Schematic representation of “Sarcomeric Remodeling” protocol (SR) and representative image showing iPSC-CMs from patient 1 & 2 and the control, immunolabelled with cardiac troponin (green), lamin B1 (grey) and DAPI (magenta) at day 25. Scale bar: 50µm. (b) Left panel: Quantification showing the distribution of cells based on the presence or absence of cardiac troponin in patient’s and control iPSC-derived cells. Two-way Analysis of Variance with Sidak’s multiple comparison test (N=4, independent differentiations with a minimum of 300 cells per condition); p-value shown as 0.0322 (*). The error bars represent mean ± SEM. Right Panel: Quantification showing the distribution of striated vs non-striated cardiac troponin-positive cells in the patient and the control group. Two-way Analysis of Variance with Sidak’s multiple comparison test (N=3, independent differentiations with a minimum of 300 cells per condition). p-value shown as 0.011(*), <0.007(**). The error bars represent mean ± SEM. (c) Representative images depicting the round and misshapen [elongated (if major axis ≥ 2.5µm), bleb, deformed] nuclei observed in patient’s hiPSC-CMs. Nuclei were immunolabelled with lamin A/C in green, lamin B1 in grey and nuclei with DAPI in magenta. Scalebar: 5µm (d) Distribution of nuclei shapes (round, elongated, Blebs, deformed or string) in cardiac troponin positive cells, striated cardiomyocytes, non-striated cardiomyocytes and cardiac troponin negative cells. (N=4, independent differentiations with a minimum of 150 nuclei per condition). Two-way analysis of variance (ANOVA) and Chi-square test with data means. p value represented as <0.0021 (**), <0.00001 (***)

Sarcomere assembly and nuclear shape are mechanically coupled through the cytoskeleton and LINC complex^16^. We investigated whether nuclear shape abnormalities were associated with defective sarcomerogenesis (Fig. 3c-d). We analyzed nuclei shape distribution within cardiomyocytes following the criteria described by Steele-Stallard *et al*.^17^ (Fig. 3c). In control iPSC-CMs, 91 ± 2% of nuclei displayed round shapes, in contrast with patient-derived iPSC-CMs that contain 66 ± 2 and 65 ± 4% round nuclei in patient-1 and -2, respectively (Fig. 3d). In addition, complete sarcomerogenesis do not impact this trend, as patient’s derived iPSC-CMs still showed a higher proportion of deformed nuclei (Fig. 3d). By contrast, in patient’s non-striated cardiomyocytes, more than 50% of nuclei were deformed (Fig. 3d). Finally, cTnT-negative cells further revealed an increase in deformed nuclei in patient-derived cells compared to control, with a notable enrichment of elongated nuclear phenotype in all iPSC lines compared to cTnT-positive cardiomyocytes (Fig. 3d). Together, these approach correlates nuclear shape abnormalities with sarcomerogenesis impairment, which appears exacerbated in *LMNA* mutant cardiomyocytes.

### *LMNA*-variant cardiomyocytes displayed reduced mitochondrial respiration and altered calcium flux

Since sarcomere organization and nuclear integrity are tightly coupled to cellular energetics^18,24^. We next investigated for altered mitochondrial respiration and identified an overall reduction in both patient lines compared to control (Fig. 4a) with noticeable variation in either mitochondria content in cells (increasing for both patients) or the amount of proteins of the oxidative phosphorylation complex (decreasing for both patients) (Fig. 4b, S4). Basal and maximal respiration were significantly decreased in patient’s cells leading to a respiratory capacity decreased to 1.7 ± 2.4 in patient-2-derived cardiomyocytes and 2.7 ± 3.1 in patient-1 compared to 3 ± 2.1 in control (Fig. 4c).

**Figure 4:**
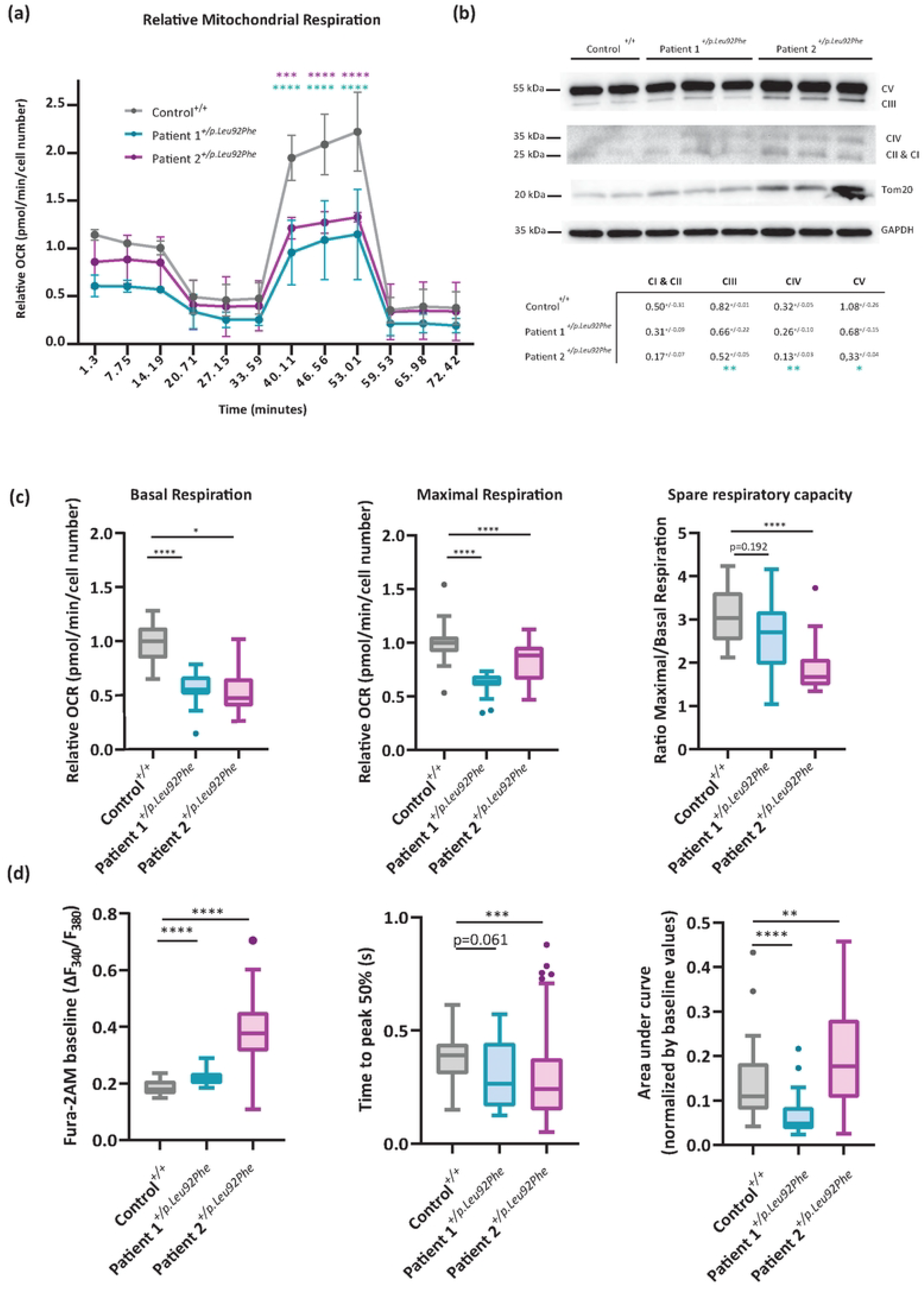
LMNA-variant iPSC cells display altered mitochondrial respiration and calcium transients. (a) Plots representing the relative oxygen consumption rate (OCR) in patients compared to control iPSC-CMs after administration of electron transport chain inhibitors (oligomycin, carbonyl cyanide-4-trifluoromethoxy phenylhydrazone (FCCP), rotenone and antimycin A). N=3 (independent differentiation), two-way analysis of variance (ANOVA) with Dunnnett’s multiple comparison test. p value represented as 0.0002 (***), 0.00001 (****) (b) Western blot analysis of OXPHOS mitochondrial electron transport chain complexes, TOM20 and GAPDH proteins expression in total extracts from patient’s cells. N=3 (independent differentiation). Table contain individual quantification of each mitochondrial electron transport chain complexes (CI&II, CIII, CIV and CV) normalized on Tom20 staining. The Student T-test was used to compare the three conditions. p value represented as <0.01 (*); <0.001 (**). (c) Tukey’s box plot showing the basal respiration, maximal respiration, and the spare respiratory capacity in patient and control iPSC-CMs. N=3 (independent differentiation). The Mann-Whitney test was used to compare the three conditions. p value represented as <0.01 (^*^), <0.0002 (^***^), <0.00001 (^****^) (d) Tukey’s box-plot representing the Fura-2AM baseline (ΔF340/F380), time to peak 50%, and area under curve in control and patient iPSC-CMs. (N=3 (independent differentiation); control iPSC-CMs = 18, patient 1 iPSC-CMs = 8, and patient 2 iPSC-CMs = 42); p value represented as <0.123 (ns), <0.0332 (^*^), <0.00001 (^****^)

Given the close interdependence between mitochondrial function and calcium handling^19^, we next analyzed calcium flux in patient’s cells that show a significant increase of the baseline calcium levels, associated with nearly half reduction in the time to peak in both patient cell lines (Fig. 4c, S3d). Additionally, the area under curve (AUC) appears quite different, with either a reduction for patient-1 or an increase for patient-2. To note, quantification of the intracellular pumps SERCA2, located in the sarcoplasmic reticula, did not reveal noticeable expression variation (Fig. S3e). Altogether, our data highlight that the *LMNA*:c.274C>T variant is also correlated with calcium homeostasis disruption that could contribute to impaired contractility in patient-derived iPSC-CMs.

## Discussion

In this study, we examined the cellular effects of the *LMNA* p.Leu92Phe variant, located in a region critical for lamin dimerization and filament assembly^20^. Using patient-derived induced pluripotent stem cell-derived cardiomyocytes, we linked this variant to altered nuclear architecture during structural remodeling, with noticeable changes in mitochondrial function and calcium handling.

Consistent with *LMNA*-associated cardiomyopathies^21,22^, *LMNA* p.Leu92Phe iPSC-derived cardiomyocytes displayed nuclear abnormalities, indicative of defective filament assembly and protein misfolding^20,23^. These defects weaken nuclear lamina integrity, reducing resistance to mechanical stress and leading to nuclear deformation^24^. Our data further suggest impaired cytoskeletal remodeling, a key process in cardiomyocyte maturation^25^. Recent work shows that sarcomeres grow through tension-driven division, which depends on stable mechanical coupling between the cytoskeleton and nucleus^26^. Under cytoskeletal remodeling conditions, control cells formed organized, striated contractile structures, whereas *LMNA*-mutant cells showed disorganized sarcomeres and pronounced nuclear collapse. A defective lamina likely disrupts the mechanical signaling required for sarcomere adaptation, explaining the late-onset disease: development proceeds normally, but long-term mechanical stress leads to progressive failure and dilated cardiomyopathy.

Metabolic analysis revealed reduced mitochondrial respiration, including mitochondria content alterations. Given the link between calcium signaling and mitochondrial function^19^, these defects likely relate to the altered calcium handling observed, forming a feedback loop that could exacerbate contractile dysfunction.

Several limitations should be noted. First, the use of related patient-derived lines introduces shared genetic background, preventing full exclusion of modifying factors. To note, our patients are from different sex, possibly highlighting sex-linked modifiers of *LMNA*-associated cardiomyopathy. Validation in unrelated cohorts or CRISPR-generated isogenic controls will be necessary. Second, while our approach identifies structural and metabolic consequences under mechanical stress, it does not resolve the precise signaling pathways connecting nuclear defects to impaired remodeling; further transcriptomic or temporal analyses will be needed. Finally, although this variant has been associated with lipodystrophy, our cohort shows a cardiac phenotype. The simplified 2D *in vitro* model cannot fully capture systemic influences that may determine tissue-specific disease manifestation.

Overall, our findings link *LMNA* p.Leu92Phe to defects in nuclear structure, sarcomere remodeling, calcium handling, and mitochondrial function, providing a basis for contractile dysfunction in *LMNA* related cardiomyopathy.

## Materials and methods

### Study approval

The Investigation Committee of the Hospital Civil of Lyon (HCL) including members of the National Reference Center for Inherited Arrhythmias of Lyon, Department of Cardiac Electrophysiology approved the clinical evaluation. Studies are conformed with the principles outlined in the Declaration of Helsinki. Subjects were being fully informed about the research, potential risks and benefits, and how data and samples would be used. The written consent has been obtained from all participants.

### Generation of human induced pluripotent stem cell lines

The iPSC lines were generated from dermal biopsies. Fibroblasts were reprogrammed by Sendai viruses (CytoTune^™^-IPS 2.0 Sendai Reprogramming kit, Life Technologies). The iPSC clones were expanded on mouse embryonic fibroblasts (MEFs) feeder cells in KSR-FGF2 medium (DMEM/F12 supplemented with 0.1% β-mercaptoethanol, 20% knockout serum replacement, 10 ng/mL basic fibroblast growth factor, 2 mmol/L l-glutamine and 1% NEAA). Until P10, colonies were mechanically passaged with a needle and adapted to feeder-free culture conditions: stem cell-qualified Matrigel-coated plates (0.1 mg/mL; BD Bioscience) with iPS Brew (Miltenyi Biotec).

### Mycoplasma detection

Mycoplasma detection was performed using the MycoAlert™ kit (LONZA, LT07-318).

### Culturing of hiPSCs

hiPSCs were cultured on Matrigel® coated dishes with StemMACS PSC-Brew XF medium (Miltenyi Biotec, cat. 130-127-865) and cultured at 37 °C, 5% CO_2_. Cells were passaged at 70-80% confluency using Accutase (Innovative Cell Technologies) or Gentle Cell Dissociation Reagent (GCDR; STEMCELL Technologies, cat. 100-0485). HiPSCs were cryopreserved in StemMACS™ CryoBrew (Miltenyi Biotec, cat. 130-109-558) and stored in liquid nitrogen.

### Cardiomyocyte Differentiation

Differentiation was performed using the StemMACS™ CardioDiff Kit XF, human (Miltenyi Biotec, cat. 130-125-125). Briefly, hiPSCs were dissociated with Accutase® for 3 min at 37 °C, pelleted (200 × g, 4 min, 20°C), resuspended in Mesoderm Induction medium (MIm) and seeded at 4.10^5^ cells/cm^2^. Cells were cultured in MIm for 24h (Day 0-1), followed by a switch to Cardiac Cultivation medium (CCm) for 24 hours (Day1-2) and then into Cardiac Induction medium (CIm) for 24 hours (Day 2-3). Cells were maintained in CCm for 15 days, dissociated using TrypLE™ Express (Thermo Fisher Scientific) and replated from day 16 to 19 under metabolic selection in a lactate-based medium. HiPSC-CMs were then replated in CCm on Matrigel-coated Ibidi® chamber slides.

### Immunocytochemistry

Cells were fixed in 4% paraformaldehyde containing 4% sucrose for 20 min at room temperature. Following PBS washes, cells were permeabilized with 0.5% Triton X-100 for 10 min and blocked in 1% BSA for 30 min. Primary antibodies were diluted in 1% BSA and applied overnight at 4 °C. Incubation with secondary antibodies was performed for 1 h at room temperature. Antibodies are listed in Table S2.

### Western blot

Samples were loaded onto 12 or 4-15% acrylamide gels and migrated at 130V for 90 min. Wet blotting system (BioRad) was used to transfer proteins to PVDF membranes (Millipore, Immobilon-P, IPVH00010). Membranes were then saturated in 5% BSA in 0.1% Tween20 – 1X TBS for 1h at RT and were incubated in primary antibodies overnight at 4°C. Following washes, membranes were incubated in HRP conjugated secondary antibodies in 1% BSA in TBST for 1h at RT. Following washes, detections of proteins were carried out using clarity Western ECL substrate (BioRad, 1705060) and ChemiDoc imaging system (BioRad) (Fig. S4). Antibodies are listed in Table S2.

### Flow Cytometry

Dissociated cells were fixed in 3% PFA for 15 min, washed, permeabilized (PBS + 1% BSA + 0.1% saponin), and stained with anti-cTnT-FITC antibody (Cardiac troponin; Miltenyi Biotec, cat. 130-119-674; 2 µL/tube, 45 min, ice). Cells were washed, resuspended in 0.5 mM EDTA-PBS, and analyzed on a MACSQuant10 cytometer.

### Calcium Flux Assay

hiPSC-CMs were replated on Matrigel® coated coverslips for ≥5 days, loaded with 2 µM Fura-2 AM (Thermo Fisher Scientific, cat. F1221) for 30 min at 37 °C in the dark. After washes, ratiometric calcium imaging was performed using the IonOptix acquisition system^8^.

### Seahorse XF Mito Stress Test

HiPSC-CMs were dissociated with TrypLE Express, seeded at 40,000-60,000 cells/well on Matrigel-coated 96-well XF plates (Agilent), and cultured 3-4 days in maturation medium. Cells were incubated 1 h without CO_2_ at 37°C in assay medium (bicarbonate-free DMEM with 25 mM glucose, 1 mM sodium pyruvate, and GlutaMAX), and sequential injection of oligomycin (1 µM), FCCP (2 µM), and rotenone + antimycin A (1 µM each) was performed.

### Genomic DNA Extraction and Sanger Sequencing

Genomic DNA extraction was performed using the PureLink™ Genomic DNA Mini Kit (Invitrogen, cat. K182001). Exon 1 of *LMNA* was amplified with GoTaq® G2 Hot Start polymerase (Promega, cat. M7401) using forward primer 5’-ACT CCG AGC AGT CTC TGT CCT-3’ and reverse primer 5’-GCA AAG TTA TCG GCC TCC AGG-3’. PCR products were resolved on a 1% agarose gel, visualized under UV illumination, and sequenced to confirm the genotype (Fig. S2a).

### Cell lines authentication by Short Tandem Repeat (STR) genotyping

The amplification was performed by Polymerase chain reaction with AmpFlSTR® Identifiler® Plus kit (Applied Biosystems). The amplified products were run on 3500 DNA Analyzer (Applied Biosystems). Data generated was analyzed using GeneMapper® Software version 4.0 (Applied Biosystems) (Table S3).

### *In silico* protein homology analysis

Sequence alignment of Lamin A/C across species was carried out using CADD, REVEL an PolyPhen2 (Fig. S1a).

### Statistical analysis

For comparison of the groups, data were assessed for normality using the Shapiro-Wilk test and presented using GraphPad Prism version 8.4 (Dotmatics). The data is represented as mean with standard error mean (SEM) if the data is normally distributed and median with interquartile range (IQR) if not normally distributed. Tests used are mentioned in the figure legends.

### Experimental Rigor and Replication Standards

To safeguard against batch-to-batch variability, we established a structured working cell bank for all iPSC lines. Throughout this study, we define an **independent biological replicate (N)** as an entirely separate experimental run originating from fresh cryovial, which was independently expanded, maintained and differentiated. For all single-cell morphometric and imaging analyses, we tracked a minimum of 50 distinct nuclei or cells per condition within each differentiation batch.

### Statement for Artificial Intelligence (AI)

The authors did not use generative AI or AI-assisted technologies in the development of this manuscript.

## Data availability

The datasets produced in this study are available in the following databases: https://www.ebi.ac.uk/biostudies/studies/S-BSST2333

## Declaration of interests

The authors declare no competing interests.

## Data and code availability

Primary data are available upon reasonable request to the corresponding author (V.G).

## Acknowledgements

This work was funded by the ATIP-AVENIR Program and VetAgro Sup Grant. We acknowledge the Platform iPSC of Nantes and particularly Dr. Laurent David as well as Dr. Julien Courchet from INMG-PGNM of Lyon.

## Notes

### Competing Interest Statement

The authors have declared no competing interest.

